# Timecourse of corticospinal excitability for observed action: evidence of early suppression followed by return to baseline without facilitation

**DOI:** 10.64898/2026.06.09.731129

**Authors:** Baptiste M. Waltzing, Marcos Moreno-Verdú, Elise E. Van Caenegem, Laurine F. Boidequin, Charlène Truong, Raphael Hamel, Robert M. Hardwick

**Affiliations:** Institute of Neurosciences, UCLouvain, Belgium Avenue Mounier 54, 1200, Bruxelles, Belgium; School of Sport, Exercise, and Rehabilitation Sciences, University of Birmingham

## Abstract

**Introduction:** Action observation modulates corticospinal excitability, with most previous studies indicating an increase in excitability in the muscles involved in the observed movement. In addition, previous work suggests that modulation of corticospinal excitability could be specific to the timing of the stimulation, muscle, and direction of movement. Here we examined the influence of these factors on corticospinal excitability.

**Method:** Participants observed stimuli presenting a static hand, followed by an image of the endpoint of an index/little finger abduction movement. Transcranial magnetic stimulation was delivered at time points from 100-800ms after movement onset. Stimuli were presented in various orientations to study possible effects of anatomical positioning and movement direction, compared relative to the control condition of a static hand.

**Results:** Corticospinal excitability was lower at early timings (100-400ms), before rising to a plateau at later timings (500-800ms) which did not differ from the static hand condition. This facilitation was muscle-specific, with higher excitability for the muscle involved in the observed movement. By contrast, the relative direction of movement did not influence corticospinal excitability.

**Discussion:** These results replicate the time-dependent modulation of corticospinal excitability induced by action observation; however, we argue that simply interpreting such effects as an increase in excitability may be overly simplistic. In line with previous studies, we argue that the choice of control condition used during action observation studies may be critical to the overall direction of effects.

## Introduction

Action Observation is known to modulate corticospinal excitability (Fadiga et al., 1995). Previous studies have typically shown that there is an increase in corticospinal activity specifically for the muscles involved in the observed movement (Fadiga et al., 1995; Urgesi et al., 2006). For example, when observing an abduction movement of the index finger, the amplitude of motor evoked potentials (MEPs) recorded in the agonist muscle, the first dorsal interosseous (FDI), increases specifically compared to that measured in an uninvolved hand muscle such as the abductor digiti minimi (ADM). However, the increase in corticospinal activity during action observation must remain below a certain threshold to avoid the production of unwanted overt imitations; it has therefore been proposed that an inhibitory mechanism could take place either in parallel during action observation and/or be triggered when excitability reaches a certain threshold, to prevent imitation (Naish et al., 2014). This hypothesis is supported by several studies that have shown a decrease in corticospinal excitability rather than an increase during the observation of actions (Sartori et al., 2012; Arias et al., 2014). The difference observed between these studies could be explained by the fact that there is a fine balance between inhibition and excitation that could be influenced by several factors, such as the observer’s intention (Hardwick et al., 2012), instruction to not move during the task (Villiger et al., 2011), motivation (Cheng et al., 2007), the baseline of comparison used (for a review see (Naish et al., 2014)), or the timing of observation (Arias et al., 2014; Lepage et al., 2010). As action observation is also used in the medical field for rehabilitation of movement related disorders (Ciullo et al., 2026) better characterising the effects of these factors could improve the effectiveness of such neurorehabilitation protocols. Further studies are therefore needed to investigate the influence of these factors on the effects of action observation.

As mentioned above, the balance between inhibition and excitation during action observation is likely time-dependent (Arias et al., 2014; Lepage et al., 2010). However, for practical simplicity, or because it was not the focus of the research question, many studies only examine a single timing (Naish et al., 2014). Furthermore, some studies do not compare the different timings studied with each other, or do not fully specify the timing (for a review see (Naish et al., 2014)); these limitations make comparisons between studies complex, and highlight once again the need for comprehensive reporting in the field of action simulation (Moreno-Verdú et al., 2024). However, some studies have nonetheless investigated multiple time points in the same task. For example, Lepage et al. (2010) showed non-specific facilitation at very early timings (60-90ms), while Cavallo et al. (2014) showed an increase in corticospinal excitability specifically for the muscles involved in the observed movement compared to those not involved, starting at 200 to 300ms. In contrast, Arias et al. (2014) showed a decrease in excitability from 360 to 500ms, and Gangitano et al. (2001) showed that excitability increased over time, with a maximum corresponding to the maximum finger aperture for a grasping task at approximately 3000ms. Based on these conflicting findings, the time course of corticospinal excitability during action observation remains to be better characterised.

In addition to these neural studies, a previous behavioural study conducted by our laboratory also suggests various modulations at early timings during action observation (Waltzing et al., 2024). Indeed, we showed that when participants had little time (300-500ms) to process a hand stimulus, an automatic response based on spatial information was given, even though it conflicted with task demands. By contrast, when participants were allowed more time to process the stimulus, they were able to produce responses based on anatomical information. This idea of two conflicting mechanisms for processing spatial and anatomical information had already been proposed by Alaerts et al. (2009). Therefore, corticospinal excitability could be modulated both in a muscle-specific manner and in a direction-specific manner depending on the type of information (anatomical and direction of movement, respectively) being processed at a given timing. However, the time course of this potential direction-specific modulation remains to be characterised at the neural level.

Our study investigated the time course of corticospinal excitability during action observation. Our hypothesis was that, as in most studies, we would observe facilitation following action observation, but the exact timing at which this modulation would occur remained uncertain. We were also interested in the time-dependent influence of direction-specific and muscle-specific modulation on corticospinal excitability. Our hypothesis was that two conflicting mechanisms would modulate corticospinal excitability: first, a direction-specific mechanism at early timings (300–500ms), and then a muscle-specific mechanism at later timings (Alaerts et al., 2009; Waltzing et al., 2024).

## Methods

### Design

The study used a within-subjects design. It was approved by the Saint-Luc - Université Catholique de Louvain hospital-faculty ethics committee (Belgium) (reference number: 30LXX111372). All procedures were reported in accordance with the ‘Action Observation’ part of the “Guidelines for Reporting Action Simulation Studies” checklist (Moreno-Verdú et al., 2024).

The study by Catmur et al. (2007), suggests a moderate effect size (d_z_ = 0.54) for muscle specific activation during action observation. A sample size calculation with G*power 3.1 for a paired two tails t-test with alpha = 0.05, power = 0.8 and a moderate effect size (d_z_ = 0.54) indicated that a sample size of more than 29 participants would be sufficient.

### Participants

A group of 34 healthy young participants were recruited into this study. Two participants were excluded due to technical issues leading to incomplete datasets. Two additional participants were excluded because more than 80% of their MEPs had to be rejected (according to the criteria detailed in Electromyography (EMG) and Transcranial Magnetic Stimulation (TMS) system); this led to less than 20 trials per condition, indicating that corticospinal excitability estimates from those two participants were unreliable (Biabani et al., 2018). This left 30 participants aged between 19 and 30 years (mean = 23.3 years, SD = 2.6 years). All participants were right-handed, had normal or corrected-to-normal vision, 26 were women and 4 were men. All participants were naive about the purpose of the experiment, provided written informed consent before the start, and were financially compensated (25€) for their participation.

### Electromyography (EMG) and TMS system

EMG was used to record the activity of the First Dorsal Interosseus (FDI) and Abductor Digiti Minimi (ADM) muscles of the right hand. The skin was first cleaned with an alcohol solution to improve conductivity. Self-adhesive pre-gelled bipolar surface electrodes (Blue Sensor N, Ambu ®, Denmark) were then placed on the belly muscles and their distal insertions in a belly-tendon montage. The reference electrode was placed on the styloid process of the ulna. The signal was sampled at 2 kHz and amplified with D360 8 Channel Amplifier (Digitimer®, England) before being online band-pass filtered (20-450 Hz) and 50-Hz notch filtered. Data were recorded for further offline analysis using the ‘Signal’ software package (version 6.04 (CED, UK)).

Monophasic TMS pulses were delivered using a Magstim 200^2^ (Magstim, Dyfed, UK) and a 40mm figure-of-eight alpha coil (Magstim Co., UK). A Visor2 neuronavigation system (ANT Neuro, The Netherlands) was used to ensure correct coil positioning throughout the experiment (Caulfield et al., 2022). The hotspot was defined as the optimal position to produce reliable MEPs for both the FDI and ADM muscles. The TMS coil handle was first positioned pointing backwards at 45º from the midline and placed tangentially over the scalp, then adjusted individually. The Resting Motor Threshold (RMT) was then determined for each participant as the lowest intensity that induced MEPs of at least 50µV from peak to peak amplitude in both muscles in at least 5 out of 10 consecutive TMS pulses (Cavallo et al., 2014). The average RMT was 50.3 ± 9.3% of maximum stimulator output. During the experiment, participants were stimulated at 120% of RMT to ensure reliable MEPs. For each trial, muscle activity before the TMS pulse was analysed to check for background noise. Trials were rejected if there was a root mean square EMG activity greater than 20 µV for the 100ms before the TMS pulse, as this could suggest voluntary muscle activation.

### Procedure

Stimuli were presented in PsychoPy2 version 2023.2.2 (Peirce et al., 2019). During the experiment, participants were comfortably seated approximately 60cm from the stimulus presentation screen with their right hand flat on a table in front of them. They were instructed to keep their right hand as relaxed and motionless as possible throughout the experiment. Before each trial, a fixation cross was presented for 2500ms. During a trial, participants first observed a hand at rest for 1100 to 1800ms, then the stimulus changed to display the endpoint of an abduction movement of the index or little finger for a total trial duration of 2900ms (Figure 1). The change from a static stimulus to a stimulus showing the endpoint of movement has shown a robust imitative effect (Catmur & Heyes, 2011; Press et al., 2005; Stürmer et al., 2000) while maintaining precise control over the timing of stimulus presentation.

**Figure 1.**
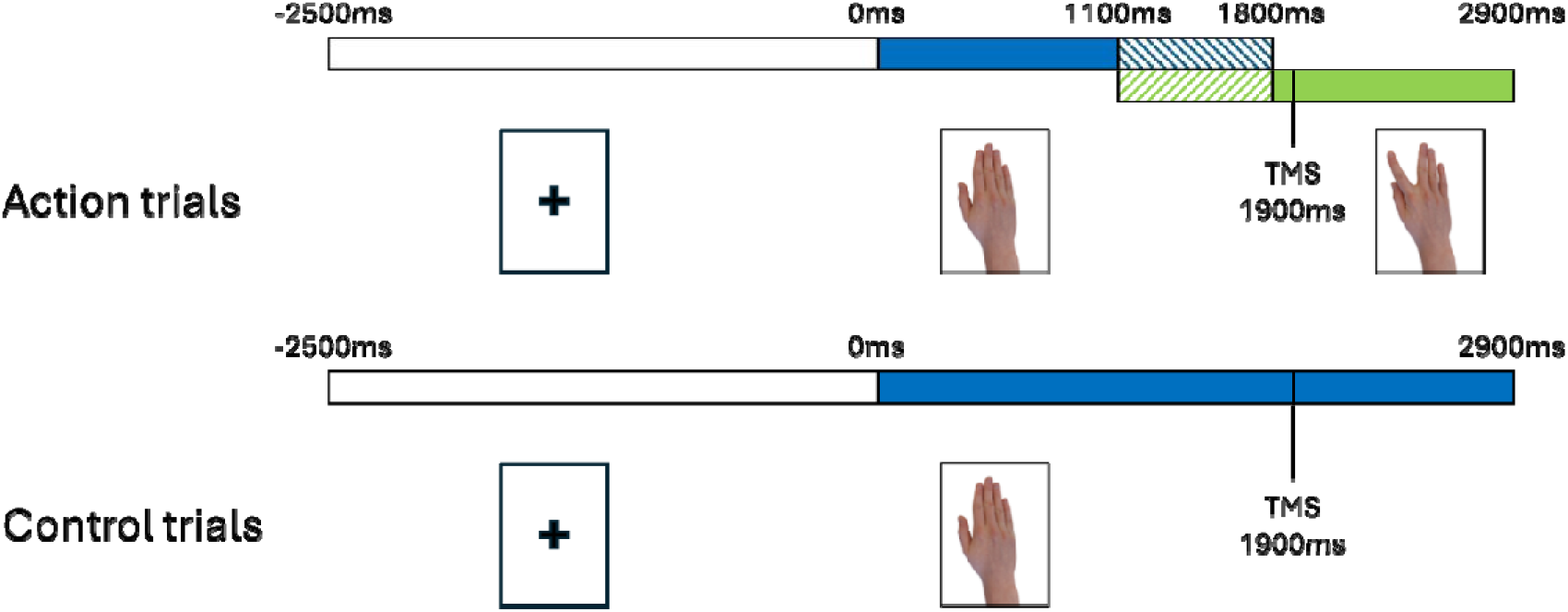
Action Trials. Before each trial, a fixation cross was presented for 2500ms. During a trial, participants first observed a hand at rest for 1100 to 1800ms (blue part). Then the stimulus changed to display the endpoint of an abduction movement of the index or little finger (green part) for a total trial duration of 2900ms. The hatched sections indicate the moment when the image could be either static or show a finger being abducted. The TMS pulse was sent 1900ms after the start of the trial, allowing stimulation to vary from 100 to 800ms after the stimulus change. *Control Trials*. For a subset of trials, the static hand was displayed for the whole duration of the trial.

The TMS pulse was always delivered at 1900ms after the start of the trial, allowing measurements ranging from 100 to 800ms in steps of 100ms after the stimulus change. For a subset of trials, the stimulus of the static hand remained for the duration of the whole trial, providing a control condition. The experiment began with a short familiarization block (10 trials) to ensure that participants understood the instructions. This was followed by 8 blocks of 60 trials, for a total of 480 trials. For the 8 stimuli with index or little finger movement, there were 6 trials for each of the 8 time points (i.e. 384 total trials with implied movement), and for the stimuli with hands at rest, there were 24 trials for each of the 4 control static hand stimuli (i.e. 96 total trials with a static hand). These numbers gave us a minimum of 24 MEPs per condition and per participant for each analysis, meeting the minimum of 20 trials recommended to have a stable MEP measurement (Biabani et al., 2018) even after trial rejection (see below). The order of trials was pseudorandomized to ensure equivalent distribution of each stimulus and timing per block throughout the experiment.

Attention checks were included to ensure that participants paid attention to the stimuli during the experiment. For each block there were 16 attention checks where participants were asked which stimulus they had seen in the previous trial. They were asked to use their left hand to press the F, G, or H buttons of a computer keyboard to indicate whether the last trial presented a stimulus involving movement of the index finger, little finger or static hand. There was no time limit for responding, and they were instructed to focus on the correct response and not on speed. The location of these attention checks during the block was randomized.

Between blocks, there was a pause (at least one minute, and longer if the participant wished). The experiment lasted a maximum of 2h.

### Stimuli

The stimuli were images of hands from a male model (Figure 2). These images could be in first- or third-person perspective, be a right or left hand, and show no movement, or an abduction of either the index or little finger. This provided a total of 12 different stimuli (2 perspectives * 2 hands * 3 finger positions). Original stimuli were created using a right hand in first-person perspective. To obtain the other stimuli, we applied transformations to the images using vertical symmetry (change in hand identity) and horizontal symmetry (change in perspective). The size of the stimulus displayed on the screen was chosen to match the actual size of the hand used as a model (approximately 17 × 9cm).

**Figure 2.**
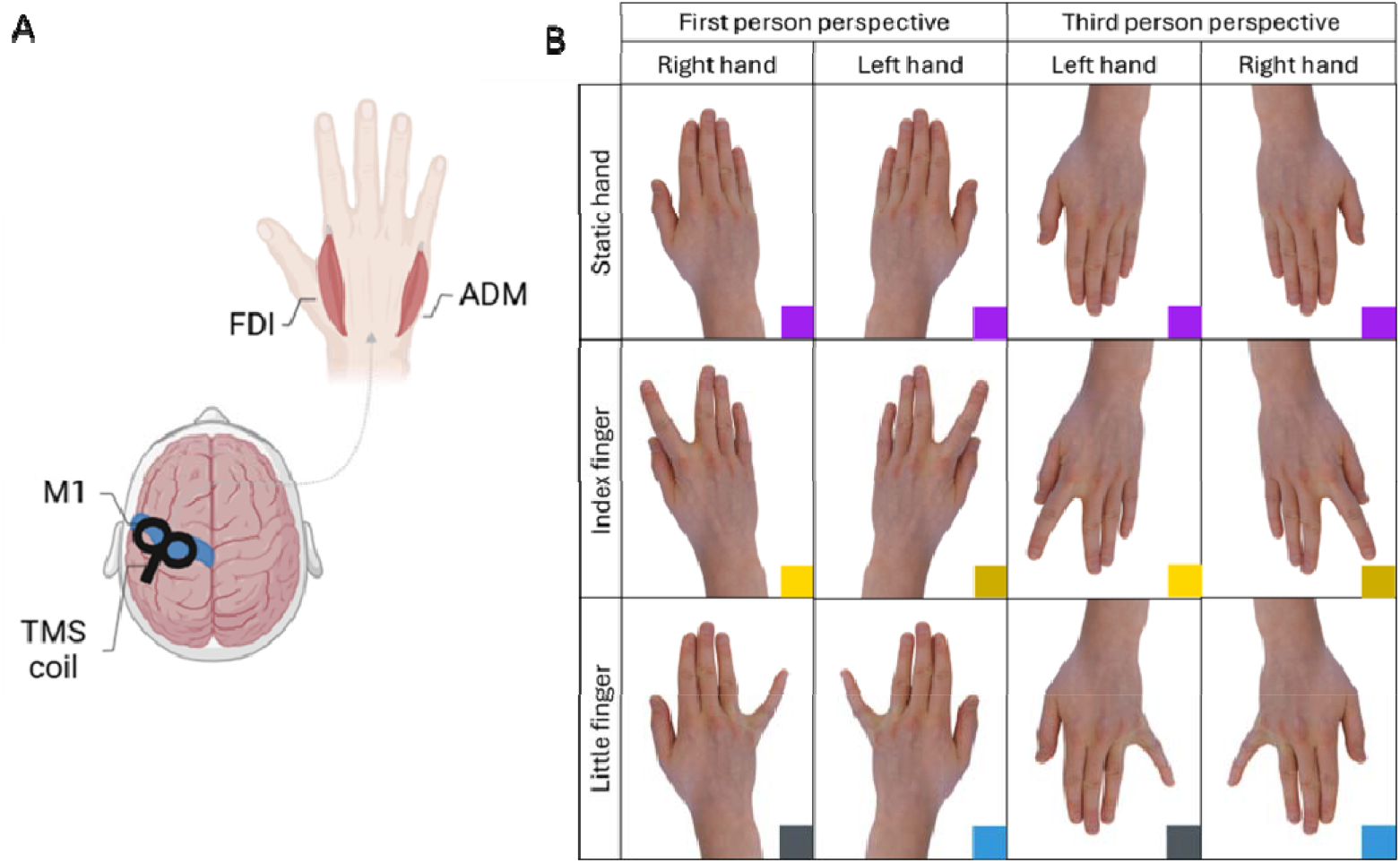
A. Illustration of the position of the TMS and the position of the participant’s hand (note the orientation is consistent with how stimuli were presented in part B). B. The 12 different stimuli used could show a right or left hand in first- or third-person perspective and an abduction movement of the index finger, little finger or static hand. The coloured squares at the bottom of the image (not shown during the experiment) represent the condition for an MEP of the FDI; purple for the static condition, yellow for anatomically congruent stimuli (stimuli showing an activation of the FDI), blue for anatomically incongruent stimuli (stimuli showing an activation of the ADM). The lighter shades are for stimuli where the direction of the movement on the screen match the direction of an abduction movement of the index finger (activation of the FDI) and the darker shades for stimuli where the direction of the movement on the screen is in the opposite direction.

**Figure 3.**
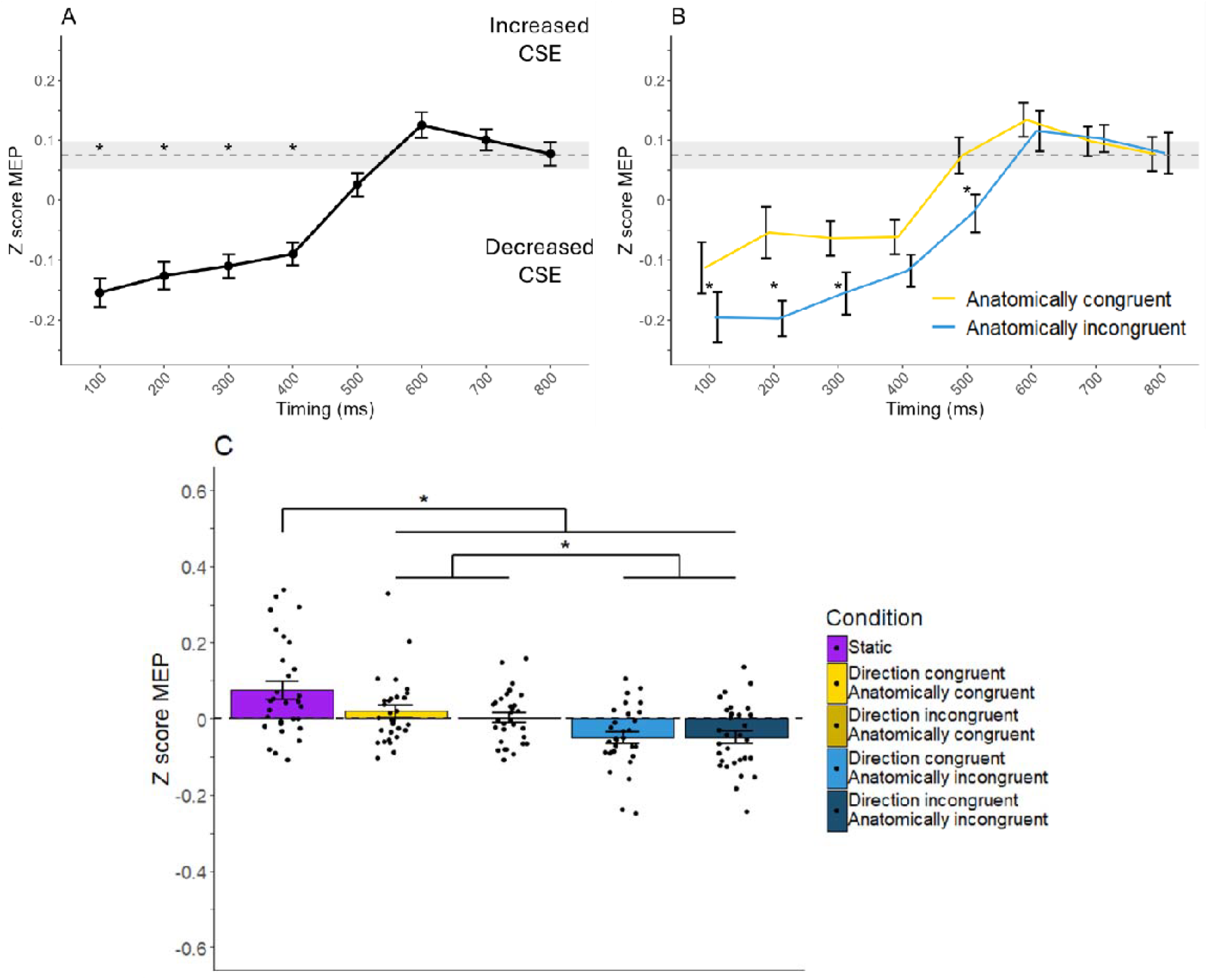
A. Main effect of timing and comparison with control condition. The dashed grey line and shaded area represent the mean and SEM for the static hand condition respectively. * In the shaded area represents a statistically significant difference with the static condition at that time point. B. Comparison between anatomically congruent and incongruent stimuli at different timings. Data from action trials were pooled across anatomic congruence. The dashed grey line and shaded area represent the mean and SEM for the static hand condition respectively. * Represents a statistically significant difference between anatomically congruent and incongruent condition at the same timing. C. Comparison of different conditions. All data were pooled together regardless of the timing. For all graphs error bars represent the standard error of the mean (SEM).

### Data analysis

All analyses were performed in R version 4.3.2 (R Core Team, 2024) and Python version 3.12. As maximal responses from the FDI and ADM are typically not equivalent, MEP amplitudes were normalised using separate z-score transformations for each muscle (i.e. for each individual participant we first pooled all measurements taken from the FDI muscle and converted them to Z-scores, then repeated this process separately for the ADM muscle). This normalization allowed us to combine MEP data from both muscles. A three-way repeated-measures ANOVA was then conducted with the within-subject factors of direction of movement (congruent vs incongruent; for examples for the FDI muscle compare light vs dark yellow colour coded stimuli in Figure 2, respectively), anatomical congruence (congruent vs. incongruent; for examples for the FDI muscle compare yellow vs blue colour coded stimuli in Figure 2, respectively), and time (8 levels: 100ms to 800ms in 100ms steps). For all analyses, Mauchly’s test was used to assess sphericity. Factors that met the assumption were analysed with uncorrected ANOVA degrees of freedom, whereas those that violated sphericity were corrected using the Greenhouse–Geisser method. Post-hoc pairwise comparisons were conducted using paired-samples t-tests where appropriate.

Control analysis was run using planned comparisons to compare action conditions with the static control condition. This analysis was conducted using paired-samples t-tests for each combination of anatomical and spatial congruence across time points. All post-hoc p-values were adjusted using the Bonferroni–Holm method (alpha = 0.05).

## Results

A significant main effect of timing was found (F=14.90, p<0.01) whereby MEPs were of lower amplitude from 100 to 400ms, and a higher amplitude from 500 to 800ms (see Table S1 for all p-values). A significant main effect of anatomical congruence (F=8.81, p<0.01) showed higher MEP amplitudes for anatomically congruent action stimuli (0.01 ± 0.06) than incongruent action stimuli (−0.05 ± 0.07) (t=2.97, p<0.01). However, the main effect of direction of movement was not significant (F=0.68, p=0.42).

Only the interaction between anatomical congruence and timing was significant (F=3.61, p<0.01). We therefore compared anatomically congruent and incongruent action stimuli for each timing and found that the two conditions were statistically different at 100-300ms and at 500ms (all t > 2.32, p < 0.03) but were not statistically different at all other timepoints examined (t=1.85, p=0.07 at 400ms and all t<0.46, p>0.65 at 600-800ms). We also compared the timings with each other separately for anatomically congruent and incongruent stimuli. For both conditions, results showed a similar pattern with lower excitability at early timings before a return to baseline. Detailed results can be found in supplementary data (Tables S2 and S3).

Control analysis showed that anatomically congruent and incongruent action stimuli both had lower MEP amplitudes than static hand stimuli (t=2.13, p=0.04 and t=3.91, p<0.01, respectively). Similarly, action stimuli between 100 and 400ms had lower MEPs amplitudes than static hand stimuli (all t>4.01, p<0.01), while MEP amplitudes of action stimuli between 500-800ms were not statistically different from those of static hand stimuli (all t<1.77, p>0.35). These results are similar when each timing is separated by anatomical congruence, with the exception of anatomically congruent stimuli at 200ms, which are not statistically different from the static hand stimuli (t=2.32, p=0.25).

## Discussion

The present study examined the time course of corticospinal excitability during action observation, as well as possible effects of congruence between the observed image and the participant’s own hand. Results demonstrated a significant effect of timing, whereby corticospinal excitability increased as presentation time lengthened. Notably, at early timings (100-400ms), MEP amplitudes for action trials were significantly *lower* than those in the static hand control condition, before rising to plateau at a level that did not differ from that of a static hand. Moreover, our results demonstrated significant modulations based on anatomical congruence with generally higher excitability for the muscles involved in the observed movement, but not the direction of movement of the stimuli.

### Effect of timing and direction of modulation

We found significant effect of the timing of stimulation relative to the presented movement; in trials presenting a movement, MEP amplitudes were relatively lower if collected shortly (100ms-400ms) after the movement, and relatively higher at later intervals (500-800ms) after the movement. However, contrary to what we had anticipated and what is generally accepted in the literature (Fadiga et al., 1995; Naish et al., 2014), at no time point did we observe higher corticospinal excitability for stimuli presenting a hand movement compared to static hand stimuli, regardless of the condition. An important point to consider is that the reference point (i.e. comparison or contrast between different conditions) used to demonstrate increased excitability varies across studies. Some authors compare two action conditions (for example, one anatomically congruent and the other not) (Catmur & Heyes, 2011). When we used this comparison, we also showed greater corticospinal excitability for our anatomically congruent condition compared to the incongruent condition for half of the time points (in the early phase after movement offset, 100-400ms). Other papers compare to a non-biological control condition such as a fixation cross (Bucchioni et al., 2013), blank screen (Ohno et al., 2011), or having the participant close their eyes (Jola et al., 2012), which does not take into account possible modulation induced by the simple observation of a hand and not by movement. For studies that compare with a static hand (Aglioti et al., 2008), as explained above, the timings are not always precisely described and are generally greater than one second after the movement (Naish et al., 2014). It is possible that if we had measured later timepoints, we would have seen an increase in excitability compared to baseline. However, this is only hypothetical and should be tested in future studies. It is also interesting to note that, unlike the study by Lepage et al. and the model proposed by Naish et al. (Lepage et al., 2010; Naish et al., 2014), we did not find a non-specific increase in excitability at our first time point (100ms). As this excitation is described as short-lived, it may have occurred prior to our first time point.

### Effect of anatomical and direction congruence

As hypothesized, we showed that there was overall higher corticospinal excitability for anatomically congruent action stimuli than for anatomically incongruent action stimuli. This result is consistent with what is generally accepted in the literature, namely that action observation increases corticospinal excitability in a muscle-specific manner (Fadiga et al., 1995; Naish et al., 2014). However, we also showed that there was an interaction between anatomical congruence and timing. In fact, there was a significant difference between the two conditions only at early timings, with higher excitability for the muscle involved in the observed movement. This again highlights the value of assessing multiple time points to capture a more detailed picture of the temporal dynamics of the effect (e.g. in the current experiment, testing at only either the 200ms or 700ms timepoint could have led to dramatically different conclusions).

In relation to the direction of the observed movement, contrary to our hypothesis, we saw no difference between directionally congruent and incongruent action conditions. This hypothesis was based on a study by Alaerts et al. (2009) where the authors suggested that corticospinal excitability could be modulated by separate mechanisms that are responsible for either muscle-specific or direction-specific effects. Our results suggest the presence of only one mechanism that induces muscle specific modulation. This may be explained by the fact that in this task, participants did not have to perform any action (no intention to imitate), unlike the behavioral task on which we based our hypothesis (Waltzing et al., 2024).

### Broader interpretation

In the TMS literature on action observation, it is generally accepted that there is an increase in corticospinal excitability during action observation. By contrast, our results, in line with those of several previous studies, show that a reduction in excitability consistent with inhibition can sometimes be observed, and that these effects are modulated by the timing of stimulus delivery. There may therefore be a fine balance between excitation and inhibition during action observation; this could vary over time and be influenced by factors such as the observer’s intention, the context, the instructions, the nature of the movement, and the choice of control/comparison condition (Loporto et al., 2011). A better understanding of these effects could also improve the applications of action observation, such as in motor learning or rehabilitation in movement-related disorders (Ciullo et al., 2026).

### Strength and limitations

One of the strengths of our paper is that we tested multiple time points and compared them with each other to obtain a clear picture of the time course of modulation. However, we did not test later timings, mainly because testing more timings in the same experiment was not practically feasible (i.e. the study was already approximately 2 hours in duration). We therefore decided to focus on the early timings that corresponded to those tested in our first behavioural study (Waltzing et al., 2024). Future studies should test later timings to obtain a more complete picture of the time course of modulation.

Another strength is that we had a sufficiently large number of participants and trials. Indeed, many previous studies were conducted on a small number of participants and sometimes with fewer trials per condition (for review see (Naish et al., 2014)) than what is now suggested to be sufficient to obtain reliable results (Biabani et al., 2018). Future studies on the subject should try to meet these same standards so that we can give equal weight to all studies.

It is also important to note that our control condition consisted of a static hand, and as mentioned previously, a wide range of different control conditions (for review see (Naish et al., 2014)) have been used in previous studies. From a practical perspective, we were not able to include additional control conditions; however, future studies could use control conditions involving a non-biological stimulus to demonstrate a potential effect specific to the observation of a body part. Such a non-biological stimulus could also display a (non-biological) movement, in order to determine whether the modulation of corticospinal excitability is not solely related to the observation of movement, or is specific to observing biological movement kinematics.

## Conclusion

Our results show that timing and anatomical congruence are factors that modulate corticospinal excitability and establish a time-course for this modulation. We also argue that the generally accepted idea that action observation induces muscle-specific facilitation should be more nuanced. Although we observed an increase in excitability over time when observing a hand action, it was never significantly higher than during the observation of a static hand. This highlights the need for further study to understand the evolution of corticospinal excitability and the potential balance between excitation and inhibition during action observation.

## Data availability statement

All data and code used for the task and data analysis can be found at: https://osf.io/urd6m/overview?view_only=25af7b1cf4534f8daa227166db040702

## Author Contributions (CRediT)

| Author | Conceptualization | Methodology | Software | Validation | Formal Analysis | Investigation | Resources | Data Curation | Writing – original draft | Writing – reviewing & editing | Visualization | Supervision | Project Administration | Funding Acquisition |
| --- | --- | --- | --- | --- | --- | --- | --- | --- | --- | --- | --- | --- | --- | --- |
| BMW | x | x | x | x | x | x | x | x | x | x | x |  |  |  |
| MMV |  |  |  |  | x |  |  |  |  | x |  |  |  |  |
| EVC |  |  |  |  |  |  |  |  |  | x |  |  |  |  |
| LFB |  |  |  |  |  |  |  |  |  | x |  |  |  |  |
| CT |  |  |  |  |  |  |  |  |  | x |  |  |  |  |
| RH |  | x |  |  |  |  |  |  |  | x |  |  |  |  |
| RMH | x | x |  | x | x |  | x |  | x | x |  | x | x | x |

## Acknowledgments

We would like to thank Julien Lambert and the members of the Neuroscience Techniques and Methods Development (NeTMeD) group for their assistance in developing the experimental setup.

## Supplementary material

**Table S1.** Post-hoc comparison between each timing. Statistically significant differences are in bold.

| Timing1 | Timing2 | statistic | df | p.adj |
| --- | --- | --- | --- | --- |
| 100 | 200 | -0.92 | 29 | 1 |
| 100 | 300 | -1.42 | 29 | 1 |
| 100 | 400 | -1.46 | 29 | 1 |
| 100 | 500 | -3.3 | 29 | **0.04** |
| 100 | 600 | -4.65 | 29 | **<0.01** |
| 100 | 700 | -5.51 | 29 | **<0.01** |
| 100 | 800 | -4.04 | 29 | **0.01** |
| 200 | 300 | -0.61 | 29 | 1 |
| 200 | 400 | -1.01 | 29 | 1 |
| 200 | 500 | -3.47 | 29 | **0.02** |
| 200 | 600 | -4.54 | 29 | **<0.01** |
| 200 | 700 | -5.27 | 29 | **<0.01** |
| 200 | 800 | -4.24 | 29 | **<0.01** |
| 300 | 400 | -0.65 | 29 | 1 |
| 300 | 500 | -3.26 | 29 | **0.04** |
| 300 | 600 | -5.05 | 29 | **<0.01** |
| 300 | 700 | -5.57 | 29 | **<0.01** |
| 300 | 800 | -4.11 | 29 | **0.01** |
| 400 | 500 | -3.6 | 29 | **0.02** |
| 400 | 600 | -5.36 | 29 | **<0.01** |
| 400 | 700 | -6.04 | 29 | **<0.01** |
| 400 | 800 | -4.04 | 29 | **0.01** |
| 500 | 600 | -2.67 | 29 | 0.15 |
| 500 | 700 | -2.45 | 29 | 0.23 |
| 500 | 800 | -1.3 | 29 | 1 |
| 600 | 700 | 0.76 | 29 | 1 |
| 600 | 800 | 1.54 | 29 | 1 |
| 700 | 800 | 0.62 | 29 | 1 |

**Figure S1.**
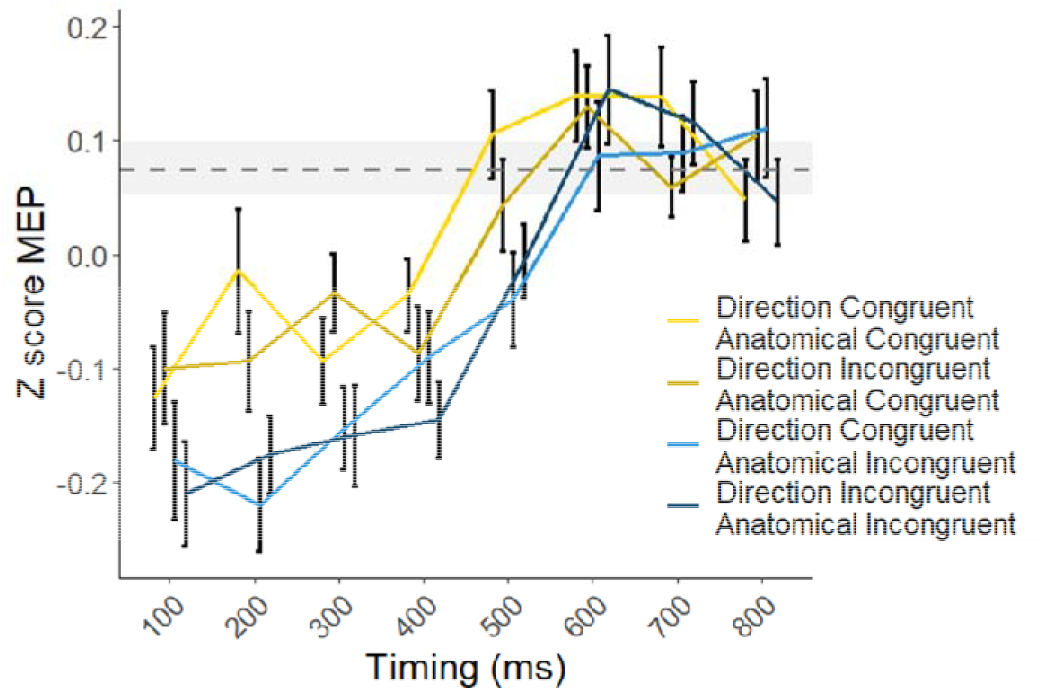
Comparison between anatomically and direction congruent and incongruent stimuli at different timings. The grey bar represents the level of excitability for the static condition and the shaded area the SEM for the static condition.

**Table S2.** Post-hoc comparison between each timing for anatomically congruent stimuli. Statistically significant differences are in bold.

| Timing1 | Timing2 | statistic | df | p.adj |
| --- | --- | --- | --- | --- |
| 100 | 200 | -1.66 | 29 | 1 |
| 100 | 300 | -1.31 | 29 | 1 |
| 100 | 400 | -1.08 | 29 | 1 |
| 100 | 500 | -3.35 | 29 | 0.05 |
| 100 | 600 | -3.96 | 29 | **0.01** |
| 100 | 700 | -4.03 | 29 | **0.01** |
| 100 | 800 | -3.02 | 29 | 0.09 |
| 200 | 300 | 0.28 | 29 | 1 |
| 200 | 400 | 0.17 | 29 | 1 |
| 200 | 500 | -2.43 | 29 | 0.30 |
| 200 | 600 | -2.93 | 29 | 0.10 |
| 200 | 700 | -2.82 | 29 | 0.13 |
| 200 | 800 | -2.37 | 29 | 0.32 |
| 300 | 400 | -0.08 | 29 | 1 |
| 300 | 500 | -3.00 | 29 | 0.09 |
| 300 | 600 | -4.39 | 29 | **<0.01** |
| 300 | 700 | -3.80 | 29 | **0.02** |
| 300 | 800 | -3.14 | 29 | 0.08 |
| 400 | 500 | -3.45 | 29 | **0.04** |
| 400 | 600 | -4.57 | 29 | **<0.01** |
| 400 | 700 | -4.46 | 29 | **<0.01** |
| 400 | 800 | -3.08 | 29 | 0.09 |
| 500 | 600 | -1.34 | 29 | 1 |
| 500 | 700 | -0.75 | 29 | 1 |
| 500 | 800 | -0.04 | 29 | 1 |
| 600 | 700 | 0.97 | 29 | 1 |
| 600 | 800 | 1.62 | 29 | 1 |
| 700 | 800 | 0.59 | 29 | 1 |

**Table S3.** Post-hoc comparison between each timing for anatomically incongruent stimuli. Statistically significant differences are in bold.

| Timing1 | Timing2 | statistic | df | p.adj |
| --- | --- | --- | --- | --- |
| 100 | 200 | 0.07 | 29 | 1 |
| 100 | 300 | -1.07 | 29 | 1 |
| 100 | 400 | -1.62 | 29 | 0.94 |
| 100 | 500 | -2.88 | 29 | 0.09 |
| 100 | 600 | -4.75 | 29 | **<0.01** |
| 100 | 700 | -6.14 | 29 | **<0.01** |
| 100 | 800 | -4.49 | 29 | **<0.01** |
| 200 | 300 | -1.43 | 29 | 1 |
| 200 | 400 | -2.16 | 29 | 0.35 |
| 200 | 500 | -4.24 | 29 | **<0.01** |
| 200 | 600 | -5.66 | 29 | **<0.01** |
| 200 | 700 | -7.13 | 29 | **<0.01** |
| 200 | 800 | -5.20 | 29 | **<0.01** |
| 300 | 400 | -1.01 | 29 | 1 |
| 300 | 500 | -3.03 | 29 | 0.07 |
| 300 | 600 | -4.86 | 29 | **<0.01** |
| 300 | 700 | -6.31 | 29 | **<0.01** |
| 300 | 800 | -4.10 | 29 | **<0.01** |
| 400 | 500 | -2.80 | 29 | 0.10 |
| 400 | 600 | -5.30 | 29 | **<0.01** |
| 400 | 700 | -5.93 | 29 | **<0.01** |
| 400 | 800 | -4.10 | 29 | **<0.01** |
| 500 | 600 | -3.53 | 29 | 0.02 |
| 500 | 700 | -3.41 | 29 | 0.03 |
| 500 | 800 | -2.27 | 29 | 0.31 |
| 600 | 700 | 0.39 | 29 | 1 |
| 600 | 800 | 1.11 | 29 | 1 |
| 700 | 800 | 0.54 | 29 | 1 |

